# *Read-SpaM*: assembly-free and alignment-free comparison of bacterial genomes with low sequencing coverage

**DOI:** 10.1101/550632

**Authors:** Anna Katharina Lau, Chris-André Leimeister, Svenja Dörrer, Christoph Bleidorn, Burkhard Morgenstern

## Abstract

In many fields of biomedical research, it is important to estimate phylogenetic distances between taxa based on low-coverage sequencing reads. Major applications are, for example, phylogeny reconstruction, species identification from small sequencing samples, or bacterial strain typing in medical diagnostics. We adapted our previously developed software program *Filtered Spaced-Word Matches (FSWM)* for alignment-free phylogeny reconstruction to work on unassembled reads; we call this implementation *Read-SpaM*. Test runs on simulated reads from bacterial genomes show that our approach can estimate phylogenetic distances with high accuracy, even for large evolutionary distances and for very low sequencing coverage. Phylogenetic trees reconstructed based on our read-based distances are largely in accordance with trusted reference trees.

**Availability:** https://github.com/burkhard-morgenstern/Read-SpaM

## 1 Introduction

Phylogeny reconstruction is a basic task in biological sequence analysis [16]. Traditionally, phylogenetic trees of species are calculated from carefully selected sets of marker genes or proteins. With the huge amount of sequencing data that are now available, genome-based phylogeny reconstruction or *phylogenomics* has become a standard approach [9, 4]. Here, the usual workflow is as follows: DNA sequencing produces a large number of reads, these reads are then assembled to obtain contigs or complete genomes. From the assembled sequences, orthologous genes are identified and multiple alignments of these genes are calculated. Finally, phylogeny methods such as *Maximum Likelihood* [45] are applied to these alignments to obtain a phylogenetic tree of the species under study. This procedure is time-consuming and error-prone, and it requires manual input from highly-specialized experts.

In recent years, a large number of alignment-free approaches to phylogeny reconstruction have been developed and applied, since these methods are much faster than traditional, alignment-based phylogenetic methods, see [51, 39, 3, 25] for recent review papers and [50] for a systematic evaluation of alignment-free software tools. Most alignment-free approaches are based on *k*-mer statistics [21, 44, 7, 48, 17], but there are also approaches based on the *length of common substrings* [47, 8, 27, 37, 32, 46], on word or spaced-word matches [38, 33, 35, 34, 1, 41] or on so-called *micro-alignments* [49, 20, 29, 28]. As has been mentioned by various authors, an additional advantage of many alignment-free methods is that they can be applied not only to complete genome sequences, but also to unassembled reads. This way, the time-consuming and unreliable genome-assembly procedure can be skipped. Assembly-free approaches can be applied, in principle, to low-coverage sequencing data. While proper genome assembly requires a coverage of around 30 reads per position, assembly-free approaches have been shown to produce good results with far lower sequencing coverage. This makes the new approach of *genome skimming* [12, 40, 10] possible, where low-coverage sequencing data are used to identify species or bacterial strains, for example in biodiversity studies [43] or in clinical applications [11, 5].

Alignment-free methods, including *Co-phylog* [49], *Mash* [35], *Simka* [2], *AAF* [14] and *Skmer* [43], have been successfully applied to unassembled reads. *Co-phylog* estimates distances using so-called *micro alignments*. In benchmark studies, *Co-phylog* could produce trees of very high quality, provided the sequencing depth was 6*X* and higher. Similarly, the programs *Mash* and *Simka* work on complete genomes as well as on unassembled reads. The required sequencing depth for these programs is comparable to the depth required by *Co-phylog*. The program *AAF* has been designed to work with unassembled data, it filters out single-copy *k-mers* to correct sequencing errors. This program produces accurate results and requires a sequencing coverage of ≥ 5 *X*.

In this paper, we introduce an alignment-free and assembly-free approach to estimate evolutionary distances, that is based on our previously introduced software *Filtered Spaced-Word Matches (FSWM)* [29]. *FSWM* is a fast program for phylogeny reconstruction that is based on gap-free local *micro-alignments*, so-called *spaced-word matches*. Originally the program has been developed to estimate distances between genome sequences; there is also an implementation of this approach called *Prot-SpaM* that can compare whole-proteome sequences to each other [28]. In this work, we adapted *FSWM* to take unassembled sequencing reads as input. Our program can compare either a set of unassembled reads from one taxon to an assembled genome of another taxon or two sets of unassembled reads to each other, each set from one taxon. Using simulated reads, we show that this method can accurately calculate distances between a complete genome and a set of reads for coverages down to 2^−5^*X*. If two sets of reads are compared, the method still works for coverages down to to 2^−2^*X*.

The paper is organized as follows: In the section 2, we shortly recapitulate how the program *FSWM* works, and we explain the modifications that we implemented to use unassembled reads as input data. In section 3, the benchmark setup and evaluation procedure are described. Section 4 describes our benchmark results, and in section 5, our results are discussed and possible future applications are mentioned.

## 2 Estimating phylogenetic distances with *FSWM* and *Read-SpaM*

For our approach, we first need to specify a binary pattern *P* of representing *match positions* and *don*’*t-care positions* [26, 22]. Let *𝓁* be the length of the pattern *P*. A spaced-word match between two DNA sequences with respect to *P* is a pair of length-*𝓁* segments, one segment from each of the sequences, such that these segments have matching nucleotides at the *match positions* of *P*. Mismatches are allowed at the *don*’*t-care* positions, see Fig. 1 for an example. In other words, a spaced-word match is a gap-free local pairwise alignment the length of *P* with matching nucleotides at the *match positions* of *P* and possible mismatches elsewhere.

**Figure 1:**
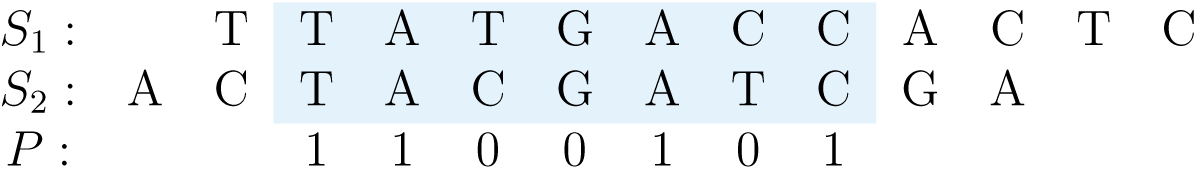
Spaced-word match between two DNA sequences *S*_1_ and *S*_2_ with respect to a binary pattern *P* = 1100101, representing *match positions* (‘1’) and *don*’*t-care positions* (‘0’). The two segments have matching nucleotides at all *match positions* of *P* but may mismatch a the *don*’*t-care* positions.

Our previously published program *FSWM* [29] estimates the *Jukes-Cantor* distance [24] between two DNA sequences as follows: first all spaced-word matches between the sequences are identified with respect to a pre-defined pattern *P*. In order to distinguish spaced-word matches representing true homologies from background spaced-word matches, a score is calculated for each spaced-word match by summing up nucleotide substitution scores for the pairs of nucleotides that are aligned at the *don*’*t-care* positions of *P*. Here we use a substitution matrix that has been proposed by Chiaromonte *et al.* [6]. Spaced-word matches with scores below some threshold value T are discarded. The remaining (‘filtered’) spaced-word matches are then used to estimate the distance between the sequences: The average number of mismatches per position is calculated for all *don*’*t-care* positions of the non-discarded spaced-word matches, and the *Jukes-Cantor* correction is used to estimate the number of substitutions per position since the sequences have evolved from their last common ancestor.

In the present study, we adapted *FSWM* to compare unassembled reads to each other or to assembled genomes. We call this implementation *Read-SpaM* (for *<u>Read</u>-based <u>Spa</u>ced-Word <u>M</u>atches*). There are two ways in which *Read-SpaM* can be used: (1) a set of unassembled sequencing reads from one taxon can be compared to a partially or fully assembled genome from another taxon; (2) a set of reads from one taxon can be compared to a set of reads from a second taxon. In both cases, all spaced-word matches between the reads and the genomes or between the reads from the first taxon and the reads from the second taxon, respectively, are identified. As in the previous version of FSWM, low-scoring spaced-word matches are discarded, and the remaining (‘filtered’) spaced-word matches are used to estimate the Jukes-Cantor distance between the two taxa. To run on short sequencing reads, we modified the length of the underlying binary patterns used in the program. While the original *FSWM* uses by default a pattern length of 112 and 12 *match positions, Read-SpaM* uses a default pattern length 72, with 12 *match positions*, i.e. with 60 *don*’*t-care positions*. As in the original *FSWM*, we are using the nucleotide substitution matrix by Chiaromonte *et al.* [6] and a threshold value of *T* = 0. That is, we discard all spaced-word matches for which the sum of the scores of the aligned nucleotides at the 60 *don*’*t-care* positions is smaller than 0. *Read-SpaM* takes *FASTA*-formatted sequence files as input, one file per input taxon.

If we want to estimate phylogenetic distances from unassembled reads as described above, we have to take sequencing errors into account. Studies have shown that *Illumina* sequencing systems have error rates of 0.24±0.06% per position [36]. Our software corrects for these errors before it calculates distances between a set of reads and a genomes, or between two different sets of reads.

## 3 Benchmark Setup

To evaluate our approach, we performed test runs on simulated reads, generated from real-world as well as from semi-artificial genome sequences. The latter sequences were obtained from real genomes by simulating evolutionary events. In both cases, the data were handled the same way. We started with pairs of full genome sequences, either real or semi-artificial. Sets of simulated reads with different coverage were then generated from these genome sequences – either from one or from both genomes in a pair of genomes – using the software tool *ART* [23]. *ART* simulates next-generation sequencing reads, it can generate reads from the three main commercial sequencing platforms with technology-specific read error models, including *Illumina*. In our test runs, we used the *Illumina HiSeq 2500* sequencing system, as it is still a widely used system in the field. Further settings were chosen as follows: The highest sequencing coverage in our study is 1*X*, and we reduced the coverage down to 2^−9^*X*, by halving the coverage in each step. This way, we cold identify a minimum coverage for which one can still obtain accurate distance estimates, for a given evolutionary distance. We used a read length of 150 bp, corresponding to the standard length of reads produced by *Illumina HiSeq 2500. ART* randomly selects positions of the genome sequences from which reads are simulated. Therefore, the generated sets of reads can vary considerably. For each pair of genomes and for each level of sequencing depth, we generated 10 sets of simulated reads.

The focus of this study is on bacterial genomes. We selected genomes of different *E. coli* strains and generated two sets of test data. First, we generated pairs of genomes consisting of a single real-world genome and a second, semi-artificial genome that we obtained by simulating nucleotide-acid substitutions, as well as insertions and deletions (indels). Indels were generated randomly with a probability of 1% at every position in the genome; the length of each indel was chosen randomly between 1 and 100, with a uniform probability distribution. Various substitution probabilities were used to generate sequence pairs with simulated evolutionary distances between 0 and around 0.75 substitutions per position. As a second set of test data, we used pairs of real-world genome sequences from different strains of *E. coli*. Generally, these genome pairs were more closely related to each other than the semi-artificial sequence pairs in the first data set. For the pairs of real-world genomes, the evolutionary distances range from 0.003 to 0.023 substitutions per position. For these pairs of real-world sequences, we used the phylogenetic distance calculated by *FSWM* from the assembled genomes as a reference, and we compared the distances obtained with *Read-SpaM* from sets of simulated sequencing reads with varying coverage.

## 4 Test Results

### 4.1 Semi-artificial pairs of genomes

For the semi-artificial sequence pairs – each pair consisting of one real genome and one artificial genome obtained from the real genome by simulated evolution –, we first applied *Read-SpaM* to estimate distances between assembled genomes and unassembled reads. Here, we generated sets of reads with sequencing coverage between 1X and 10^−9^X. As mentioned above, 10 sets of reads were generated for each genome pair and level of sequencing coverage. In Fig. 2 and 3, the average of the estimated distances is plotted against the real distance of the two genomes for distance values between 0 and *>* 0.75 substitutions per position. Standard deviations are represented as error bars.

**Figure 2:**
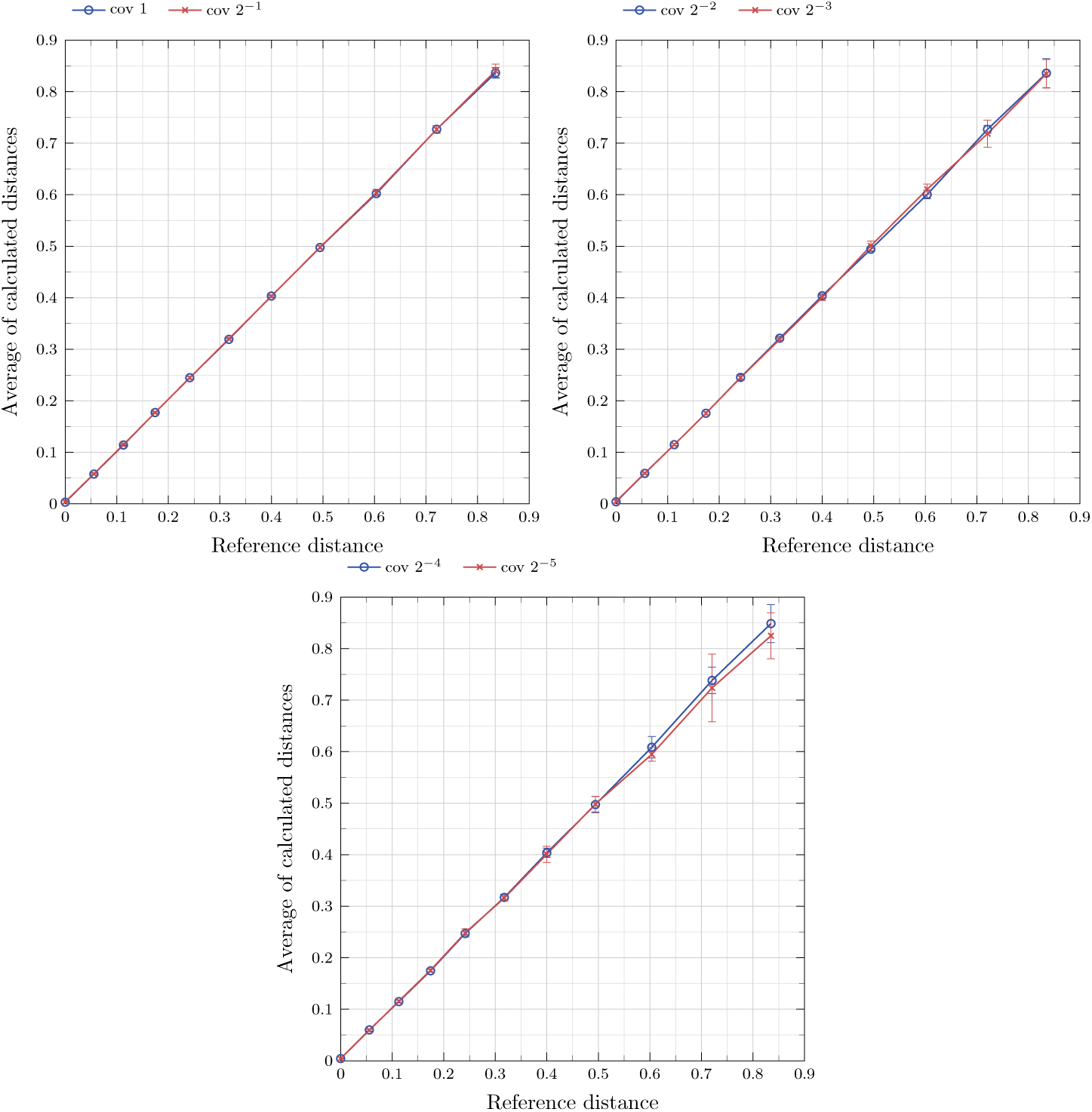
Phylogenetic distances, measured as substitutions per position, between semi-artificial assembled genomes and unassembled reads (see main text), estimated by *Read-SpaM*. Estimated distances are plotted against the real distances for sequence coverage values between 1*X* and 10^−5^*X*.

**Figure 3:**
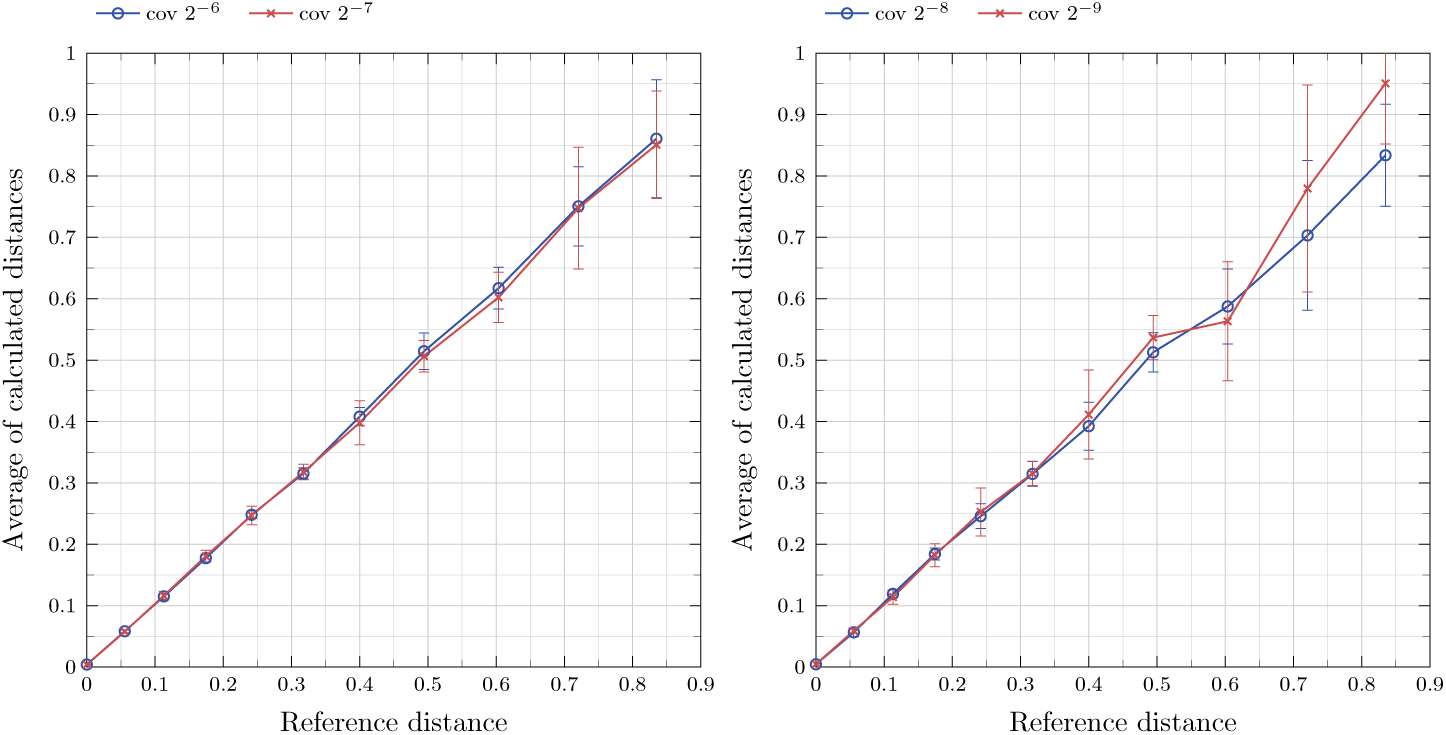
Distances between genomes and unassembled reads, plotted against the real distances as in Fig. 4, for sequencing coverage values between 2^−6^*X* and 2^−9^*X*.

Next, we used the same semi-artificial genome pairs as above, but we generated simulated reads for *both* genome sequences. The results for the comparison of unassembled reads from one genome against unassembled reads from a second genome are shown in Fig. 4. For these test data, it was not always possible to estimate distances with *Read-SpaM*, since for large evolutionary distances and/or for low sequencing coverage, no spaced-word matches with positive scores could be found. For a sequencing coverage values ≥ 2^−3^*X*, though, distances could be estimated for all data sets, i.e. up to our maximal distance of around 0.75 substitutions per position.

**Figure 4:**
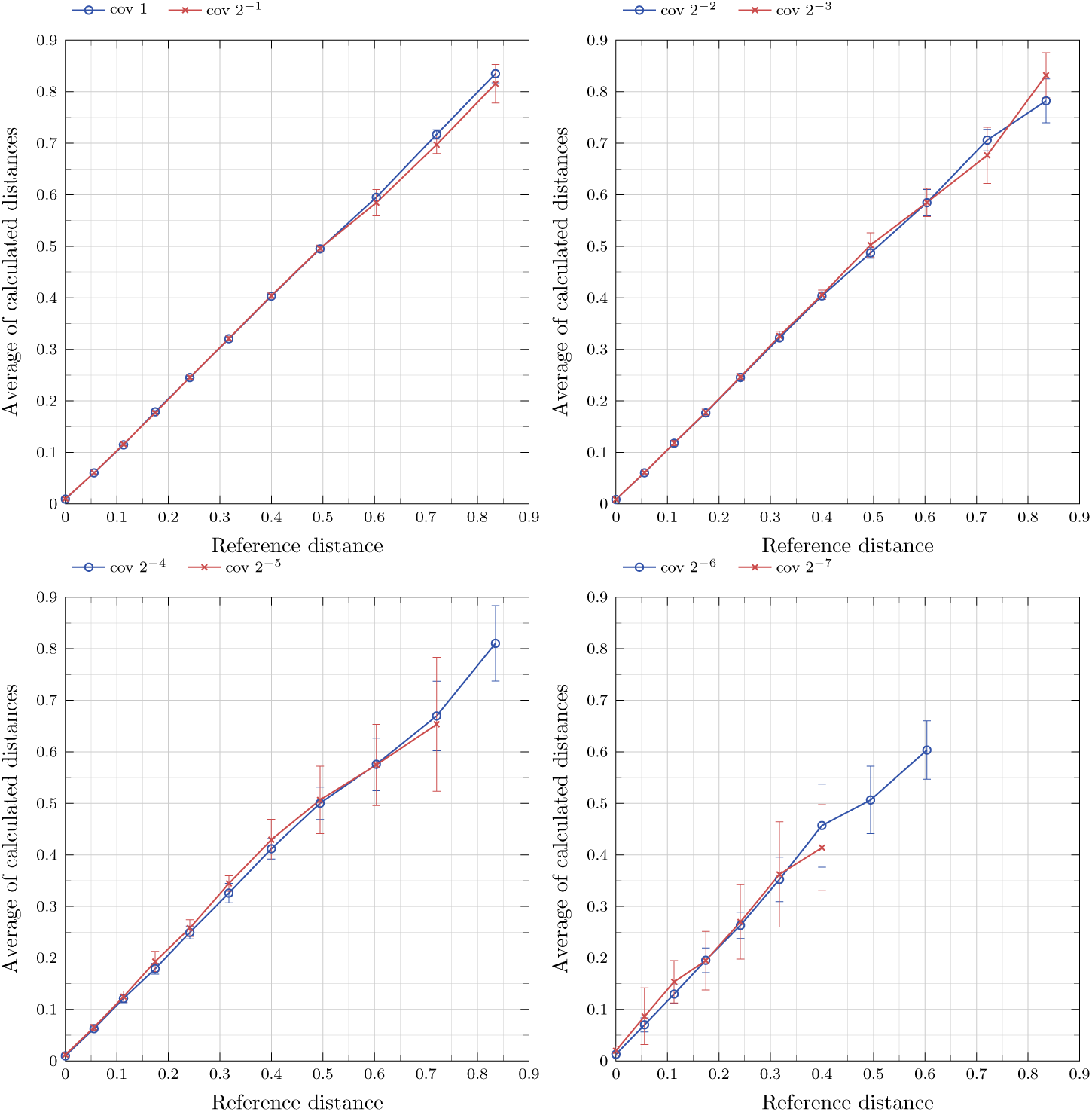
Distances estimated by *Read-SpaM* between sets of unassembled reads from two semi-artificial genomes each, plotted against the real distances, for sequencing coverage values between 1*X* and 2^−7^*X*.

### 4.2 Real-world genome pairs

In addition to the above test runs on semi-artificial genome sequences, we used pairs of real genomes from different strains of *E. coli*. We compared the distances obtained with *Read-SpaM* using unassembled reads to the distances calculated by *FSWM* calculated from the corresponding assembled genomes. Again, we first compared one assembled genome to a set of simulated reads from a second genome; then we compared a set of reads from one genome to a set of reads from another genome. The results of the test runs on real genomes are shown in Fig. 5 and 6: for each pair of genomes, distances calculated between assembled genomes and unassembled reads are shown in blue, while distances between unassembled reads from two different genomes are shown in red.

**Figure 5:**
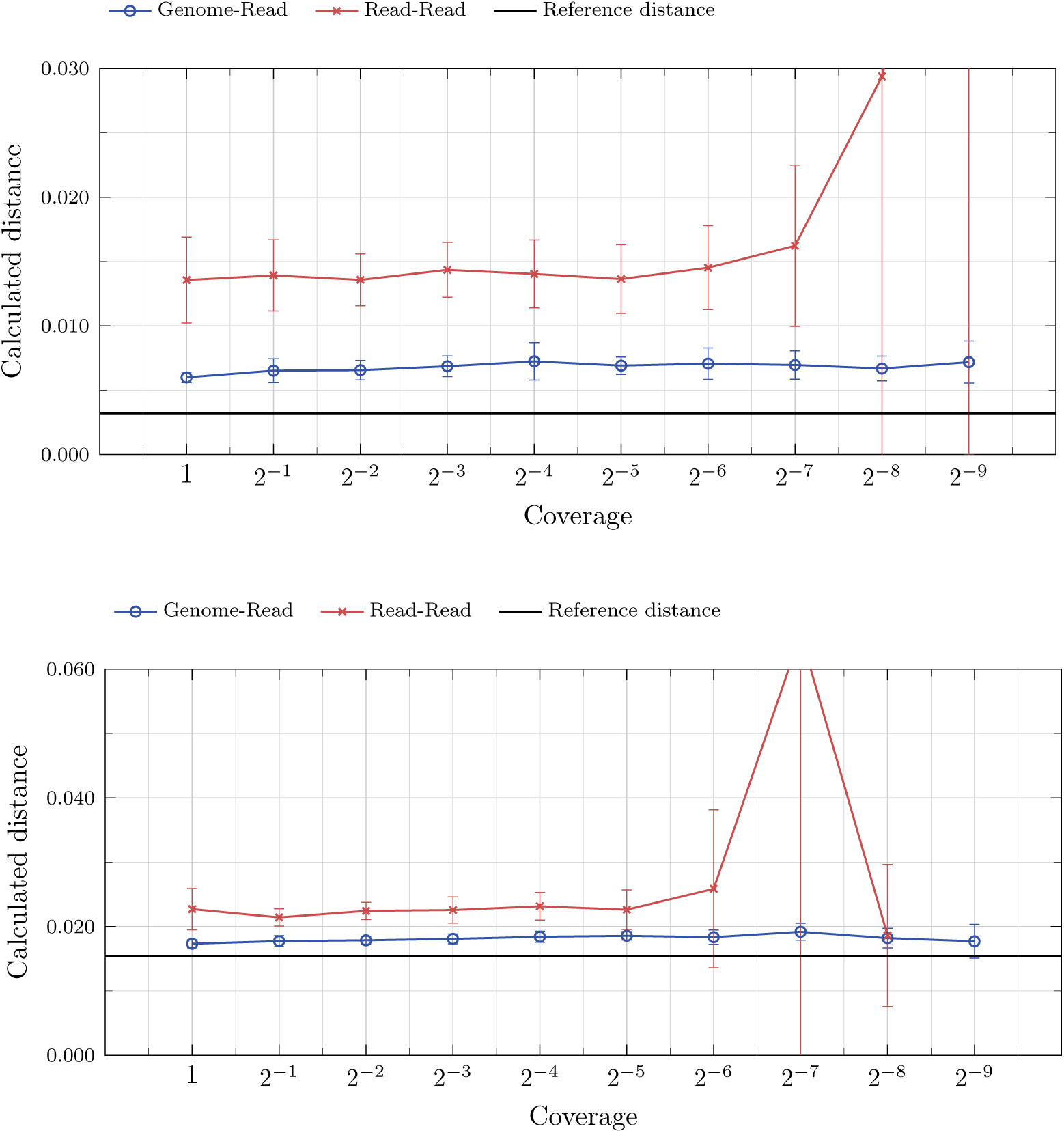
Test runs with *E. coli* strains B4Sb227 vs. BW2952 (top) and IAI1 vs. F2a2457T (bottom). As reference distances, we used the distances calculated by *FSWM* based on the assembled genomes (horizontal black lines). Distances estimated by *Read-SpaM* using unassembled simulated reads from one strain and the assembled genome from a second strain are shown in blue; distances estimated using unassembled reads from both strains are shown in red. Reads were simulated with sequencing-coverage levels from 1*X* down to 2^−9^*X*.

**Figure 6:**
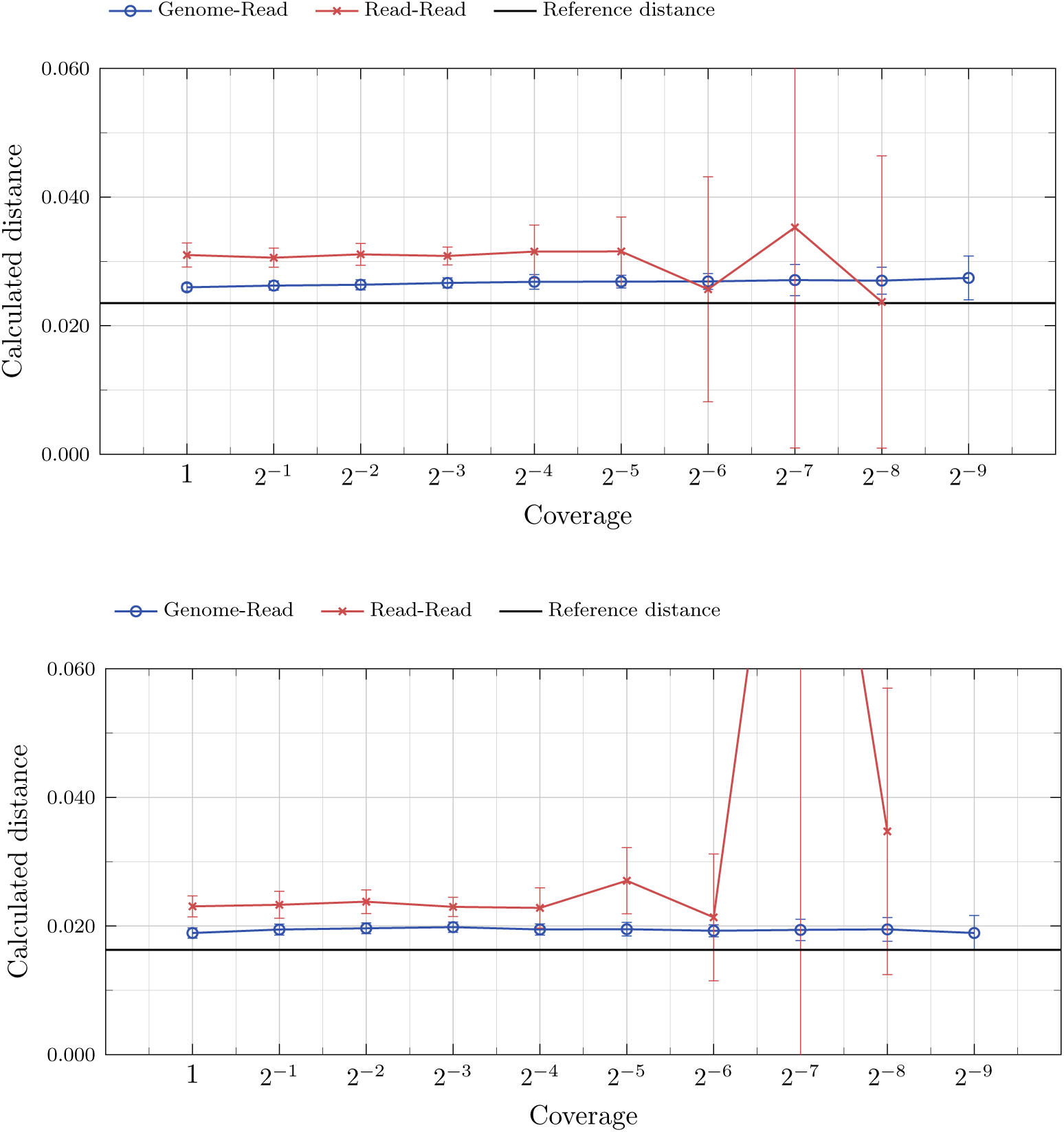
Test runs with *E. coli* ATCC8739 vs. ED1a (top) and SMS35 vs. SE11 (bottom). Blue, red and black lines as in Fig. 5

### 4.3 Wolbachia Phylogeny

Next, we applied our approach to bacterial phylogeny analysis. Here, a typical task is to find the position of a newly sequenced strain within a known tree of strains for which the assembled genome sequences are already available. As a test case, we used genome sequences of 13 *Wolbachia* strains from the lineages (“supergroups”) A - D, In addition, we used 4 strains of closely related Alphaproteobacteria as outgroup. *Wolbachia* belong to the Alphaproteobacteria and are intracellular endosymbionts of arthropods and nematodes, see [19] for classification of *Wolbachia*. As a reference tree, we used a tree published by [18].

We generated four sequence data sets, each set consisting of 12 genome sequences from different *Wolbachia* strains, 4 genome sequences from the out-group strains and a set of unassembled reads from a 13th *Wolbachia* strain taken from [18]. We then applied *Read-SpaM* to estimate phylogenetic distances within each data set, and calculated trees from these distance matrices with *Neighbor-Joining* [42] implementation from the *PHYLIP* package [15]. The resulting trees are shown in Fig. 7 and 8.

**Figure 7:**
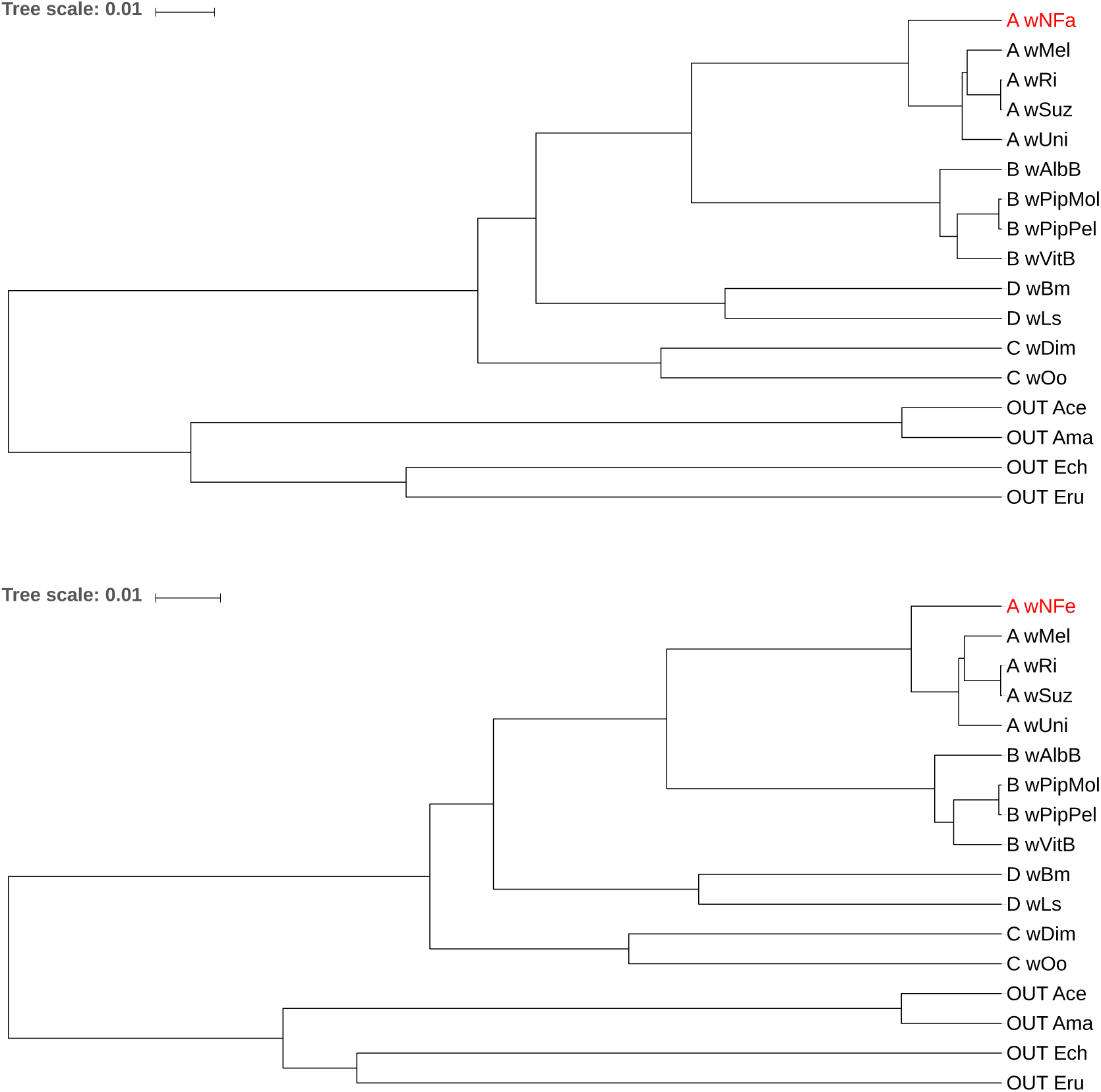
Phylogenetic trees for a set of 13 *Wolbachia* strains from super groups *A* – *D* and 4 strains from the closely related alphaproteobacterial genera Anaplasma and Ehrlichia as outgroup. For each tree, we used the full genome sequences from 12 *Wolbachia* strains and the 4 outgroup strains. For the 13th *Wolbachia* strain, we used sets of unassembled sequencing reads. The strain with the unassembled reads was *wNFa* (top) and *wNFe* (bottom).

**Figure 8:**
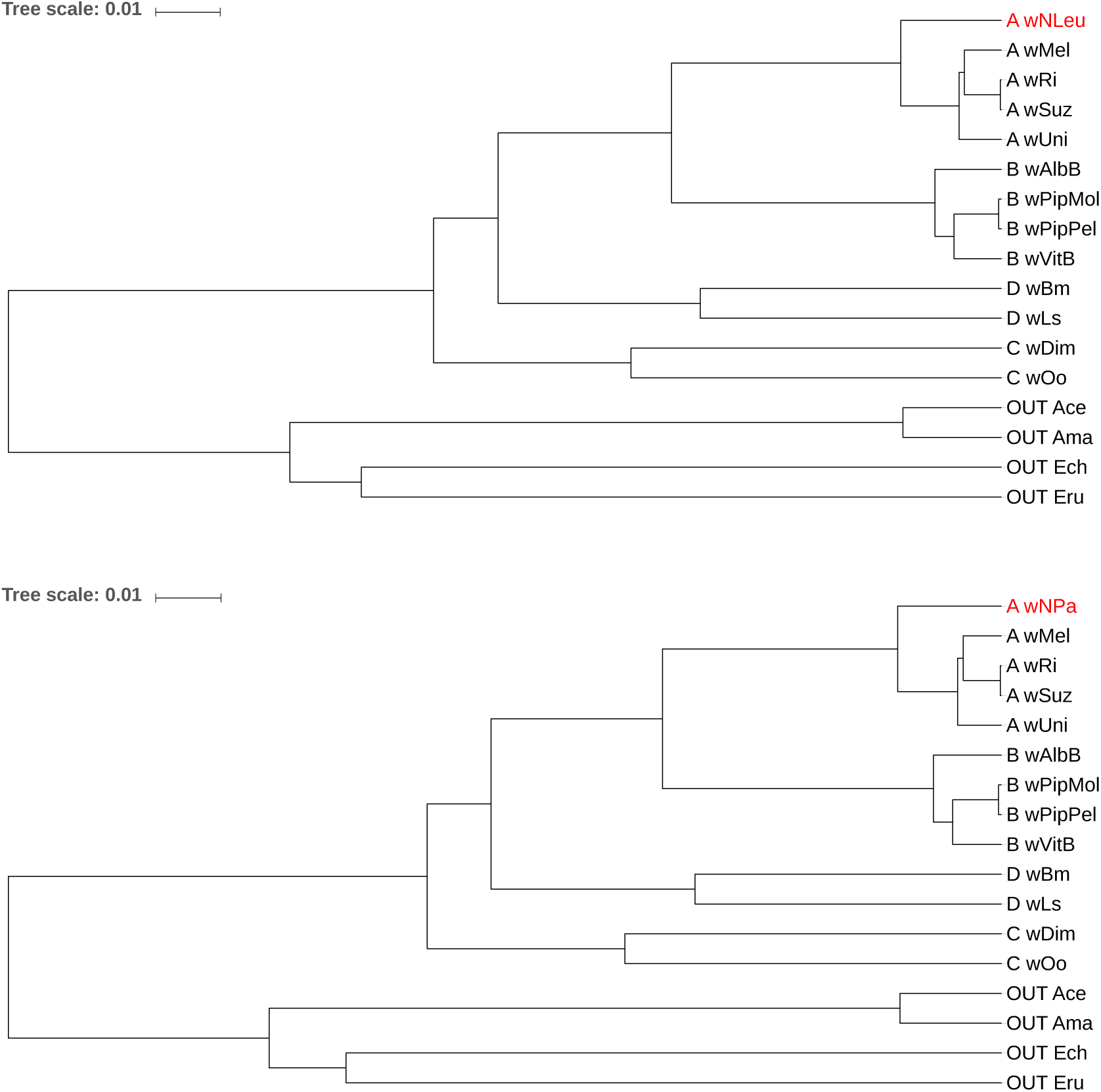
Phylogenetic trees for 17 bacterial strains, see Fig. 7. Here, we used unassembled reads from strains *wNLeu* (top) and *wNPa* (bottom) as input sequences, for the respective other strains we used their full genome sequences.

### 4.4 Runtime

As shown in Table 1. the runtime of *Read-SpaM* for comparing two strains of *E. coli* is between 0.8 s and 3.4 s, depending on the level of sequencing coverage. As a comparison, a run of *FSWM* on a data set of this size takes around 6 s. As expected, read-read comparison is faster than genome-read comparison, for each level of sequencing coverage. For both methods, the runtime decreases heavily in the beginning but only small differences can be found for a coverage below 2^−4^*X*.

**Table 1:** Runtime of *Read-SpaM* (in seconds) to estimate the distance between two strains of *E. coli*, by comparing an assembled genome to unassembled reads and by comparing unassembled reads from both strains to each other, for varying levels of sequencing coverage.

| Coverage | Genome vs. Read | Read vs. Read |
| --- | --- | --- |
| 1 | 3.404 | 2.836 |
| $2^{-1}$ | 2.426 | 1.551 |
| $2^{-2}$ | 2.024 | 1.161 |
| $2^{-3}$ | 1.857 | 0.997 |
| $2^{-4}$ | 1.737 | 0.927 |
| $2^{-5}$ | 1.755 | 0.887 |
| $2^{-6}$ | 1.641 | 0.870 |
| $2^{-7}$ | 1.637 | 0.864 |
| $2^{-8}$ | 1.662 | 0.867 |
| $2^{-9}$ | 1.628 | 0.853 |

## 5 Discussion

In this paper, we introduced *Read-SpaM*, an adaption of our previously published software *Filtered Spaced Word Matches (FSWM)* to estimate phylogenetic distances between sets of unassembled reads. We evaluated this approach on real and semi-artificial bacterial genomes with varying phylogenetic distances and for varying levels of sequencing coverage.

Fig. 2 and 3 show that, if an assembled genome from one bacterial strain is compared to unassembled reads from another strain, distances predicted by *Read-SpaM* are fairly accurate, even for very low levels of sequencing coverage and for large phylogenetic distances. For a sequencing coverage down to 2^−3^*X*, phylogenetic distances predicted by *Read-SpaM* are accurate and statistically stable for the whole range of distances that we tested, i.e. for up to around 0.8 substitutions per position. For lower levels of sequencing coverage, though, our results became less accurate and less stable for simulated strains with large evolutionary distances. But even for a coverage of 2^−9^*X* – the lowest coverage that we tested –, evolutionary distances predicted by *Read-SpaM* were quite accurate and statistically stable for distances up to around 0.5 substitutions per position.

The distance estimates between sets of unassembled reads from two taxa each are shown in Fig. 4. As can be expected, these results were not quite as accurate as estimates between reads and assembled genomes. But still, we obtained reasonably accurate and stable results for fairly large distances and low levels of sequencing coverage. For a coverage of 2^−3^*X, Read-SpaM* still produced accurate results for distances as large as 0.5 substitutions per position, while for a coverage of 2^−5^*X*, the results were still accurate for distances up to around 0.3 substitutions per position.

Our program evaluation shows that read-based estimation of phylogenetic distances with *Read-SpaM* has a high potential. The developed approach should be particularly useful for phylogenetic distances below 0.6 substitutions per position, and if unassembled reads are to be compared to assembled genomes. An important application is, for example, to search for the position of a previously unknown species in an existing phylogenetic tree, the so-called *phylogenetic placement* [31]. In this situation, low-pass sequencing can be an attractive alternative to phylogenetic barcoding based on selected *marker genes* [30, 13] to identify the phylogenetic position of an unknown species. As read-to-read comparison with *Read-Spam* still produces reliable results for sequencing coverage down to 2^−3^*X*, it is possible to estimate phylogenetic distances between strains or species for which assembled genomes are not available. We will continue to evaluate and apply our approach to further explore its potential.

## Acknowledgements

We thank Michael Gerth for discussions on the phylogeny of Wolbachia.

## Notes

#### Summary of Updates

One author, Michael Gerth, wanted to be removed from the author list, since he felt that he did not contribute enough to be listed as an author. In addition, we made slight modifications to the wording.

https://github.com/burkhard-morgenstern/Read-SpaM

## References

[1] Nathan A. Ahlgren, Jie Ren, Yang Young Lu, Jed A. Fuhrman, and Fengzhu Sun. Alignment-free 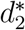 oligonucleotide frequency dissimilarity measure improves prediction of hosts from metagenomically-derived viral sequences. Nucleic Acids Research, 45:39–53, 2017.

[2] Gaëtan Benoit, Pierre Peterlongo, Mahendra Mariadassou, Erwan Drezen, Sophie Schbath, Dominique Lavenier, and Claire Lemaitre. Multiple comparative metagenomics using multiset k-mer counting. PeerJ Computer Science, 2:e94, 2016.

[3] Guillaume Bernard, Cheong Xin Chan, Yao-Ban Chan, Xin-Yi Chua, Yingnan Cong, James M. Hogan, Stefan R. Maetschke, and Mark A. Ragan. Alignment-free inference of hierarchical and reticulate phylogenomic relationships. Briefings in Bioinformatics, 22:426–435, 2019.

[4] Christoph Bleidorn. Phylogenomics. An Introduction. Springer, Berlin, 2017.

[5] Karel Břinda, Alanna Callendrello, Lauren Cowley, Themoula Charalampous, Robyn S Lee, Derek R MacFadden, Gregory Kucherov, Justin O’Grady, Michael Baym, and William P Hanage. Lineage calling can identify antibiotic resistant clones within minutes. bioRxiv, 10.1101/403204, 2018.

[6] Francesca Chiaromonte, Von Bing Yap, and Webb Miller. Scoring pairwise genomic sequence alignments. In Russ B. Altman, A. Keith Dunker, Lawrence Hunter, and Teri E. Klein, editors, Pacific Symposium on Bio-computing, pages 115–126, Lihue, Hawaii, 2002.

[7] Benny Chor, David Horn, Yaron Levy, Nick Goldman, and Tim Massingham. Genomic DNA *k*-mer spectra: models and modalities. Genome Biology, 10:R108, 2009.

[8] Matteo Comin and Davide Verzotto. Alignment-free phylogeny of whole genomes using underlying subwords. Algorithms for Molecular Biology, 7:34, 2012.

[9] Frédéric Delsuc, Henner Brinkmann, and Hervé Philippe. Phylogenomics and the reconstruction of the tree of life. Nature Reviews Genetics, 6:361–375, 2005.

[10] Dee R. Denver, Amanda M. V. Brown, Dana K. Howe, Amy B. Peetz, and Inga A. Zasada. Genome Skimming: A rapid approach to gaining diverse biological insights into multicellular pathogens. PLoS Pathogens, 12(8):e1005713, 2016.

[11] Ruud H. Deurenberg, Erik Bathoorn, Monika A. Chlebowicz, Natacha Couto, Mithila Ferdous, Silvia García-Cobos, Anna M.D. Kooistra-Smid, Erwin C. Raangs, Sigrid Rosema, Alida C.M. Veloo, Kai Zhou, Alexander W. Friedrich, and John W.A. Rossen. Application of next generation sequencing in clinical microbiology and infection prevention. Journal of Biotechnology, 243:16 –24, 2017.

[12] Steven Dodsworth. Genome skimming for next-generation biodiversity analysis. Trends in Plant Science, 20:525 – 527, 2015.

[13] Dirk Erpenbeck, Markus Steiner, Astrid Schuster, Martin J. Genner, Renata Manconi, Roberto Pronzato, Bernhard Ruthensteiner, Didier van den Spiegel, Rob W.M. van Soest, and Gert Wörheide. Minimalist barcodes for sponges: a case study classifying African freshwater Spongillida. Genome, 62:1–10, 2019.

[14] Huan Fan, Anthony R. Ives, Yann Surget-Groba, and Charles H. Cannon. An assembly and alignment-free method of phylogeny reconstruction from next-generation sequencing data. BMC Genomics, 16:522, 2015.

[15] Joseph Felsenstein. PHYLIP - Phylogeny Inference Package (Version 3.2). Cladistics, 5:164–166, 1989.

[16] Joseph Felsenstein. Inferring Phylogenies. Sinauer Associates, Sunderland, MA, USA, 2004.

[17] Umberto Ferraro-Petrillo, Gianluca Roscigno, Giuseppe Cattaneo, and Raffaele Giancarlo. Informational and linguistic analysis of large genomic sequence collections via efficient hadoop cluster algorithms. Bioinformatics, 34:18261833, 2018.

[18] Michael Gerth and Christoph Bleidorn. Comparative genomics provides a timeframe for *Wolbachia* evolution and exposes a recent biotin synthesis operon transfer. Nature Microbiology, 2:16241, 2016.

[19] Eliza Glowska, Anna Dragun-Damian, Miroslawa Dabert, and Michael Gerth. New Wolbachia supergroups detected in quill mites (Acari: Syringophilidae). Infection, Genetics and Evolution, 30:140–146, 2015.

[20] Bernhard Haubold, Fabian Klötzl, and Peter Pfaffelhuber. andi: Fast and accurate estimation of evolutionary distances between closely related genomes. Bioinformatics, 31:1169–1175, 2015.

[21] Michael Höhl, Isidore Rigoutsos, and Mark A. Ragan. Pattern-based phylogenetic distance estimation and tree reconstruction. Evolutionary Bioinformatics Online, 2:359–375, 2006.

[22] Sebastian Horwege, Sebastian Lindner, Marcus Boden, Klaus Hatje, Martin Kollmar, Chris-André Leimeister, and Burkhard Morgenstern. *Spaced words* and *kmacs*: fast alignment-free sequence comparison based on inexact word matches. Nucleic Acids Research, 42:W7–W11, 2014.

[23] Weichun Huang, Leping Li, Jason R Myers, and Gabor T Marth. ART: a next-generation sequencing read simulator. Bioinformatics, 28:593–594, 2011.

[24] Thomas H. Jukes and Charles R. Cantor. Evolution of Protein Molecules. Academy Press, New York, 1969.

[25] Gregory Kucherov. Evolution of biosequence search algorithms: a brief survey. Bioinformatics, btz272, 2019.

[26] Chris-André Leimeister, Marcus Boden, Sebastian Horwege, Sebastian Lindner, and Burkhard Morgenstern. Fast alignment-free sequence comparison using spaced-word frequencies. Bioinformatics, 30:1991–1999, 2014.

[27] Chris-André Leimeister and Burkhard Morgenstern. *kmacs*: the *k*-mismatch average common substring approach to alignment-free sequence comparison. Bioinformatics, 30:2000–2008, 2014.

[28] Chris-Andre Leimeister, Jendrik Schellhorn, Svenja Dörrer, Michael Gerth, Christoph Bleidorn, and Burkhard Morgenstern. Prot-SpaM: Fast alignment-free phylogeny reconstruction based on whole-proteome sequences. GigaScience, 8:giy148, 2019.

[29] Chris-André Leimeister, Salma Sohrabi-Jahromi, and Burkhard Morgenstern. Fast and accurate phylogeny reconstruction using filtered spaced-word matches. Bioinformatics, 33:971–979, 2017.

[30] Xiwen Li, Yang Yang, Robert J. Henry, Maurizio Rossetto, Yitao Wang, and Shilin Chen. Plant DNA barcoding: from gene to genome. Biological Reviews, 90:157–166, 2015.

[31] Benjamin Linard, Krister Swenson, and Fabio Pardi. Rapid alignment-free phylogenetic identification of metagenomic sequences. Bioinformatics, btz068, 2019.

[32] Burkhard Morgenstern, Svenja Schöbel, and Chris-André Leimeister. Phylogeny reconstruction based on the length distribution of *k*-mismatch common substrings. Algorithms for Molecular Biology, 12:27, 2017.

[33] Burkhard Morgenstern, Bingyao Zhu, Sebastian Horwege, and Chris-André Leimeister. Estimating evolutionary distances between genomic sequences from spaced-word matches. Algorithms for Molecular Biology, 10:5, 2015.

[34] Kevin D. Murray, Christfried Webers, Cheng Soon Ong, Justin Borevitz, and Norman Warthmann. kWIP: The k-mer weighted inner product, a de novo estimator of genetic similarity. PLOS Computational Biology, 13:e1005727, 2017.

[35] Brian D Ondov, Todd J Treangen, Páll Melsted, Adam B Mallonee, Nicholas H Bergman, Sergey Koren, and Adam M Phillippy. Mash: fast genome and metagenome distance estimation using minhash. Genome Biology, 17:132, 2016.

[36] Franziska Pfeiffer, Carsten Gröber, Michael Blank, Kristian Händler, Marc Beyer, Joachim L Schultze, and Günter Mayer. Systematic evaluation of error rates and causes in short samples in next-generation sequencing. Scientific Reports, 8:10950, 2018.

[37] Cinzia Pizzi. MissMax: alignment-free sequence comparison with mismatches through filtering and heuristics. Algorithms for Molecular Biology, 11:6, 2016.

[38] Gesine Reinert, David Chew, Fengzhu Sun, and Michael S. Waterman. Alignment-free sequence comparison (I): Statistics and power. Journal of Computational Biology, 16:1615–1634, 2009.

[39] Jie Ren, Xin Bai, Yang Young Lu, Kujin Tang, Ying Wang, Gesine Reinert, and Fengzhu Sun. Alignment-free sequence analysis and applications. Annual Review of Biomedical Data Science, 1:93–114, 2018.

[40] Sandy Richter, Francine Schwarz, Lars Hering, Markus Böggemann, and Christoph Bleidorn. The utility of genome skimming for phylogenomic analyses as demonstrated for glycerid relationships (Annelida, Glyceridae). Genome Biology and Evolution, 7:3443–3462, 2015.

[41] Sophie Röhling, Thomas Dencker, and Burkhard Morgenstern. The number of *k*-mer matches between two DNA sequences as a function of *k*. bioRxiv, doi:10.1101/527515, 2019.

[42] Naruya Saitou and Masatoshi Nei. The neighbor-joining method: a new method for reconstructing phylogenetic trees. Molecular Biology and Evolution, 4:406–425, 1987.

[43] Shahab Sarmashghi, Kristine Bohmann, M. Thomas P. Gilbert, Vineet Bafna, and Siavash Mirarab. Skmer: assembly-free and alignment-free sample identification using genome skims. Genome Biology, 20:34, 2019.

[44] Gregory E. Sims, Se-Ran Jun, Guohong A. Wu, and Sung-Hou Kim. Alignment-free genome comparison with feature frequency profiles (FFP) and optimal resolutions. Proceedings of the National Academy of Sciences, 106:2677–2682, 2009.

[45] Alexandros Stamatakis. RAxML-VI-HPC: maximum likelihood-based phylogenetic analyses with thousands of taxa and mixed models. Bioinformatics, 22:2688–2690, 2006.

[46] Sharma V. Thankachan, Sriram P. Chockalingam, Yongchao Liu, and Ambujam Krishnan Srinivas Aluru. A greedy alignment-free distance estimator for phylogenetic inference. BMC Bioinformatics, 18:238, 2017.

[47] Igor Ulitsky, David Burstein, Tamir Tuller, and Benny Chor. The average common substring approach to phylogenomic reconstruction. Journal of Computational Biology, 13:336–350, 2006.

[48] Susana Vinga, Alexandra M. Carvalho, Alexandre P. Francisco, Luís M. S. Russo, and Jonas S. Almeida. Pattern matching through Chaos Game Representation: bridging numerical and discrete data structures for biological sequence analysis. Algorithms for Molecular Biology, 7:10, 2012.

[49] Huiguang Yi and Li Jin. Co-phylog: an assembly-free phylogenomic approach for closely related organisms. Nucleic Acids Research, 41:e75, 2013.

[50] Andrzej Zielezinski, Hani Z Girgis, Guillaume Bernard, Chris-Andre Leimeister, Kujin Tang, Thomas Dencker, Anna Katharina Lau, Sophie Röhling, JaeJin Choi, Michael S Waterman, Matteo Comin, Sung-Hou Kim, Susana Vinga, Jonas S Almeida, Cheong Xin Chan, Benjamin James, Fengzhu Sun, Burkhard Morgenstern, and Wojciech M Karlowski. Benchmarking of alignment-free sequence comparison methods. bioRxiv, 10.1101/611137, 2019.

[51] Andrzej Zielezinski, Susana Vinga, Jonas Almeida, and Wojciech M. Karlowski. Alignment-free sequence comparison: benefits, applications, and tools. Genome Biology, 18:186, 2017.

